# Ifit2 regulates murine-coronavirus spread to the spinal cord white matter and its associated myelin pathology

**DOI:** 10.1101/2022.06.06.494972

**Authors:** Madhav Sharma, Debanjana Chakravarty, Ajay Zalavadia, Amy Burrows, Patricia Rayman, Nikhil Sharma, Lawrence C. Kenyon, Cornelia Bergmann, Ganes C. Sen, Jayasri Das Sarma

**Affiliations:** Department of Biological Sciences, Indian Institute of Science Education and Research Kolkata, Mohanpur, West Bengal, India; Department of Inflammation and Immunity, Lerner Research Institute, Cleveland Clinic, Ohio, USA; Department of Pathology, Anatomy and Cell Biology, Thomas Jefferson University, Philadelphia, Pennsylvania, USA; Department of Neurosciences, Cleveland Clinic, Ohio, USA

**Keywords:** Ifit2, coronavirus, demyelination

## Abstract

Ifit2, an interferon-induced protein with tetratricopeptide repeats 2, plays a critical role in restricting neurotropic murine β-coronavirus RSA59 infection. RSA59 intracranial injection of Ifit2 deficient (-/-) compared to wild type (WT) mice results in impaired acute microglial activation, associated with reduced CX3CR1 expression, which consecutively limits migration of peripheral lymphocytes into the brain, leading to impaired virus control followed by severe morbidity and mortality. While the protective role of Ifit2 is established for acute viral encephalitis, less is known about its influence on demyelination during the chronic phase of RSA59 infection. Our current study demonstrates that Ifit2 deficiency causes extensive RSA59 viral spread throughout both the spinal cord grey and white matter and is associated with impaired CD4**^+^** T cell infiltration. Cervical lymph nodes of RSA59 infected Ifit2**^-/-^** mice showed reduced activation of CD4**^+^** T cells and impaired IFNγ expression during acute encephalomyelitis. Furthermore, blood-brain-barrier integrity was preserved in the absence of Ifit2 as evidenced by integral, tight junction protein ZO-1 expression surrounding the meninges and blood vessels and decreased Texas red dye uptake. In contrast to WT mice exhibiting only sparse myelin loss, the chronic disease phase in Ifit2**^-/-^** mice was associated with severe demyelination and persistent viral load, even at low infection doses. Overall, our study highlights that Ifit2 provides antiviral functions by promoting acute neuroinflammation and thereby aiding virus control and limiting severe demyelination.

**Author Summary:** The role of interferons in providing protective immunity against viral spread and pathogenesis is well known. Interferons execute their function by inducing certain genes collectively called as interferon stimulated genes (ISGs) among which Interferon-induced protein with tetratricopeptide repeats 2, Ifit2, is known for restricting neurotropic viral replication and spread in the brain. So far, not much has been investigated about its role in viral spread to the spinal cord and its associated myelin pathology. Towards this our study using neurotropic murine-β-coronavirus and Ifit2 deficient mice demonstrate that Ifit2 deficiency causes extensive viral spread throughout grey and white matter of spinal cord accompanied by impaired microglial activation and CD4^+^ T cell infiltration. Furthermore, infected Ifit2 deficient mice showed impaired activation of T cells in cervical lymph node and Blood-Brain-Barrier was relatively intact. Ifit2 deficient mice developed viral induced severe chronic neuroinflammatory demyelination accompanied by the presence of ameboid shaped phagocytotic microglia/macrophages.

## Introduction

The immunomodulatory properties of interferons make them useful in the treatment of multiple sclerosis (MS) which is a chronic inflammatory neurodegenerative demyelinating disease of the central nervous system (CNS)[1–3]. The action of interferons is mediated by the expression of numerous genes called Interferon stimulated genes (ISGs) that encode for antiviral and immunomodulatory factors[4]. Among these, interferon-induced protein with tetratricopeptide repeat 2 (Ifit2) is a restriction factor against Rabies-virus, Vesicular-stomatitis-virus, West-nile-virus, Sendai-virus and, murine β-coronavirus, Mouse hepatitis virus (MHV)[5–10]. Several studies, including those with MHV, have explored underlying antiviral regulatory mechanisms of Ifit2[5, 11]. Intracranial infection with the demyelinating strain MHV-A59 or its spike protein isogenic recombinant strain RSA59 initiates activation of innate immune responses followed by prominent adaptive immunity which controls infectious virus below the detection limit by day 15 Post-infection (p.i.). However, viral RNA persists at a very low level. A gradual increase in myelin pathology with or without axonal loss is evident as early as day 7 p.i. and reaches its peak during the persisting phase (day 30 p.i.), mimicking certain pathological features of MS[12–16]. Intracranial inoculation of Ifit2**^-/-^** mice with low doses of RSA59, which only elicits mild symptoms in wildtype (WT) mice, caused pronounced morbidity and mortality accompanied by uncontrolled virus replication with significantly impaired microglial activation, reduced expression of CX3CR1, and reduced recruitment of NK1.1**^+^** and CD4**^+^** T cells into the brain[11]. While the role of Ifit2 in acute inflammation is well established, the impact of Ifit2 deficiency on virus spread to the spinal cord and its associated neuroinflammatory demyelination remains to be investigated. The current study reveals significantly heightened, indiscriminate viral spread within spinal cord grey and white matter, reduced Iba1**^+^** microglia/macrophages, and impaired local CD4**^+^** T cell infiltration in Ifit2**^-/-^** spinal cords, similar to that observed in the brains during acute infection. Reduced activation of CD4**^+^** T cells in draining CLN revealed a contribution of peripheral immune dysregulation to loss of T cell function. Despite elevated virus load, BBB integrity was maintained in Ifit2**^-/-^** compared to WT mice. Although Ifit2**^-/-^** mice given a comparatively low doses of RSA59 survived, they developed severe progressive clinical symptoms associated with augmented white matter demyelination as well as grey matter pallor compared to sparse demyelination not affecting grey matter in WT mice. Demyelinated lesions in Ifit2**^-/-^** mice exhibited significantly more ameboid phagocytic microglia/macrophages. Moreover, viral antigen was more abundant during chronic disease relative to the sparse detection in WT spinal cords. The data imply that elevated viral persistence associated with ongoing detrimental microglia/macrophage activity may amplify severe chronic progressive demyelination in Ifit2**^-/-^** mice.

## Results

### Ifit2 deficiency significantly increased RSA59 spread but restricted microglial activation in the spinal cords at the acute phase of neuroinflammation

Inoculation of WT mice with RSA59 into the brain near the lateral geniculate nuclei results in rapid viral spread to the olfactory bulb, cerebral cortex, ventral striatum/basal forebrain, hippocampal region, midbrain, medulla, followed by infection of the brainstem and deep cerebellar white matter and ultimately, the spinal cord white matter[16]. Previous studies showed that Ifit2 played a significant antiviral role against RSA59 dissemination within the brain in 4-5-week-old mice even at inoculation doses of 2000 PFUs, which is 1/10th of half of the LD50 dose. Ifit2-/- mice developed severe clinical distress and hind limb paralysis compared to WT mice and succumbed to infection by day 8 p.i.[11]. In the current study, RSA59 replicates profusely throughout the spinal cord grey and white matter in Ifit2**^-/-^** mice (Fig. 1B). In contrast, viral spread in RSA59 infected WT mice was mainly contained to the white matter of the dorsal columns with very little spillover into the grey matter as shown by immunohistochemical detection of nucleocapsid protein at day 5 p.i., representing the enhanced virus replication (Fig. 1A, 1C). In contrast to the profuse viral spread, microglial activation was significantly impaired in RSA59 infected Ifit2**^-/-^** spinal cords, as evidenced by limited Iba1 expression at day 5 p.i. (Fig. 1 D-F). No significant differences were observed in astrocytic GFAP expression in the spinal cords of RSA59 infected Ifit2**^-/-^** compared to WT mice (Fig. 1G-I). Thus, Ifit2 deficiency restricts microglial/macrophage activation throughout the CNS upon acute RSA59 infection.

**Fig. 1:**
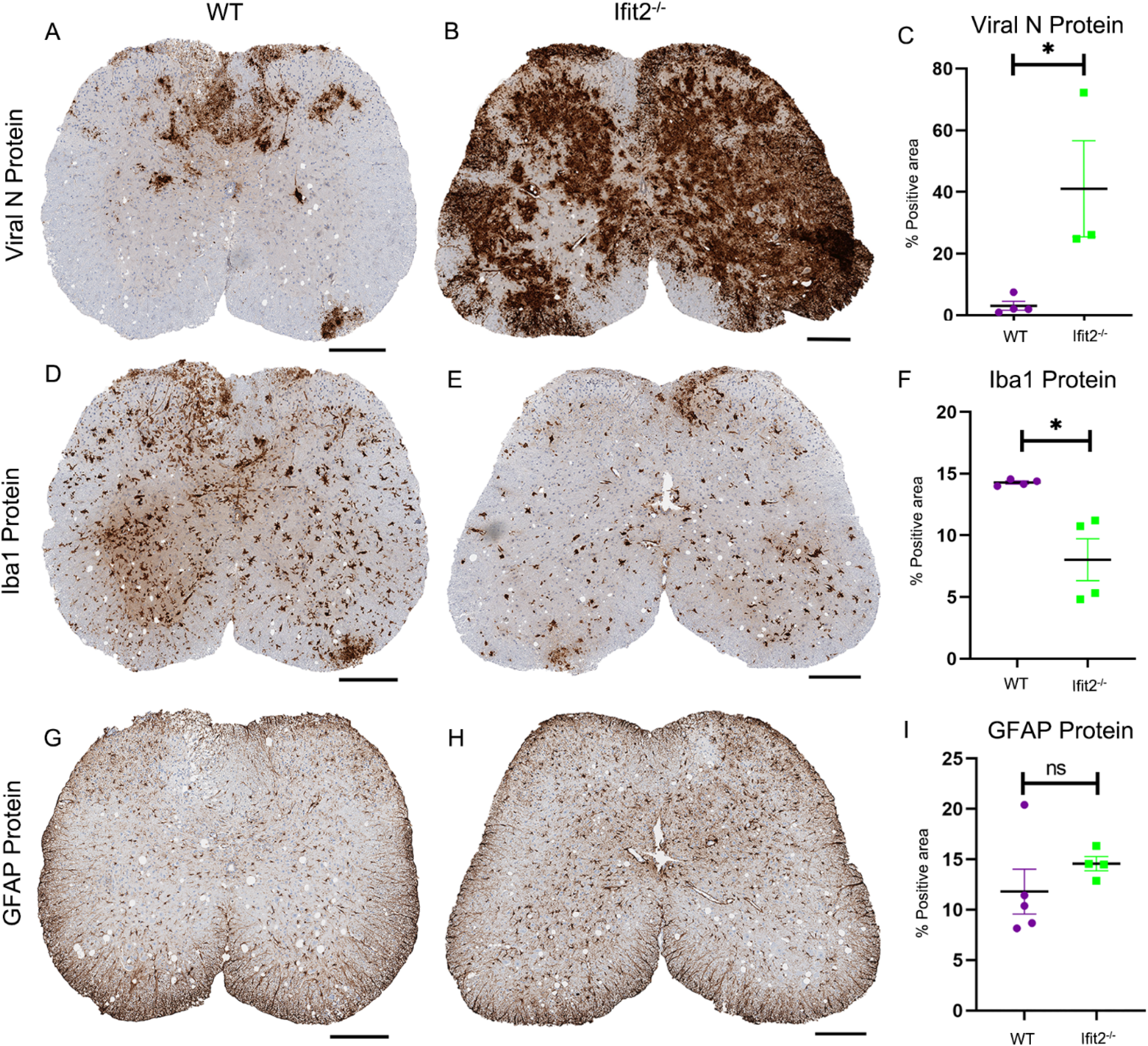
Ifit2 deficiency increases virus spread but restricts microglial activation in RSA59 infected spinal cords during acute infection. 5μm thick serial sections from spinal cord tissues at day 5 p.i. were stained for viral nucleocapsid protein (Fig. 1A, 1B), Iba1 for microglia/monocyte/macrophages (Fig. 1D, 1E) and GFAP for astrocytes (Fig. 1G,1H). Quantification of viral N protein, Iba1 and GFAP expression is graphically represented in Fig. 1C, 1F, and 1I respectively. The scale bar is 200µm. The experiment was repeated thrice, N=3 with 4-5 mice each time. Asterix (*) represents differences that are statistically significant by Student’s unpaired t-test analysis. The error bars represent SEM

### Ifit2 deficiency reduces CD4^+^ leukocyte infiltration in RSA59 infected acutely inflamed spinal cords

Activated leukocytes infiltrate the brain of RSA59 infected mice prior to clearance of infectious virus from the CNS[11, 17]. Neutrophils are the first cells to enter the brain, followed by circulating monocytes and T lymphocytes. Flow cytometric analysis of spinal cord cells was performed to assess differences in the infiltration of specific leukocyte populations in RSA59 infected WT versus Ifit2**^-/-^** mice at days 3, 5, and 7 p.i. Gating on total CD45**^+^** cells allowed distinction between CD45**^lo/in^** microglia and CD45**^hi^** peripheral infiltrating leukocytes. The CD45**^hi^** population increased markedly at days 5 and 7 p.i. in both WT and Ifit2**^-/-^** infected mice (Fig. 2B). However, CD45**^hi^** cells were significantly reduced in Ifit2**^-/-^** infected mice compared to WT infected mice at day 7 p.i. (Fig. 2A,2C). To assess whether reduced leukocyte recruitment to the spinal cord involved a specific cell type, we monitored early infiltrating CD45**^hi^** cells via immunophenotyping. Spinal cord-derived cells were assessed for CD4**^+^** T and CD8**^+^** T cell subsets (Fig. 2D). The number of CD4**^+^** T cells increased between days 5 to 7 p.i. and days 3 to 5 p.i. in WT and Ifit2**^-/-^** mice respectively (Fig. 2E) whereas CD8**^+^** T cell population increased between days 3 to 5, days 5 to 7 and days 3 to 5 p.i. in WT and Ifit2**^-/-^** mice respectively (Fig 2G). Both WT and Ifit2**^-/-^** spinal cords harbored similar lymphocyte cell numbers at days 3 and 5 p.i, however, there were fewer CD4**^+^** T cells in the spinal cord of Ifit2**^-/-^** compared to WT mice at day 7 p.i. (Fig. 2F, 2H). CD4**^+^** T cell interaction with microglia/macrophages is required for effective viral antigen clearance and maintenance of CNS homeostasis[17]. Staining for Ly6G to mark neutrophils also revealed no significant changes between the groups throughout days 3 to 7 p.i (Fig. 2I-K). Thus, impaired CD4**^+^** T cell and microglia/macrophage communication may underlie the disease severity in Ifit2**^-/-^** mice.

**Fig. 2:**
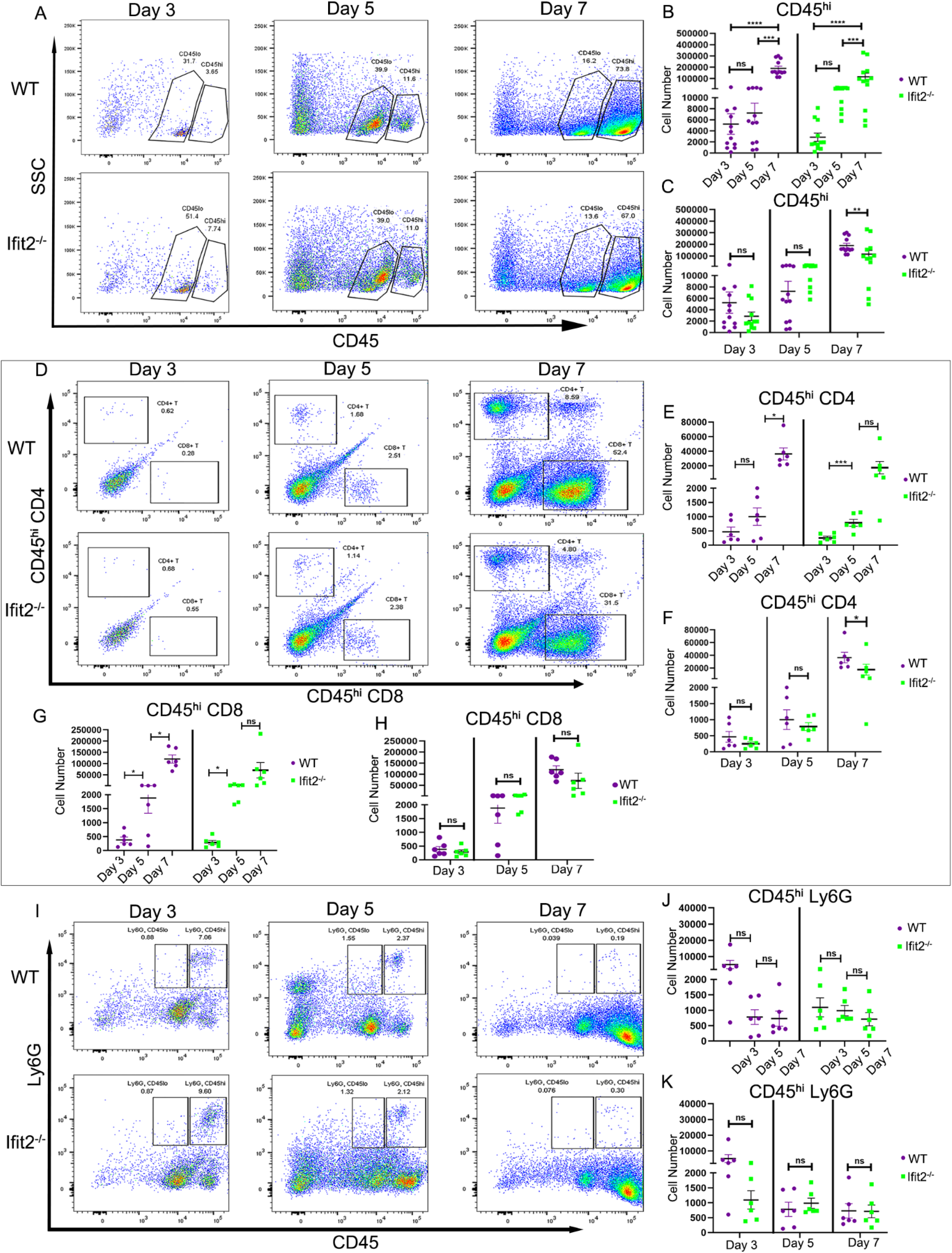
Ifit2 deficiency reduces CD4+ T cell infiltration into RSA59 infected spinal cords. Spinal cords from infected WT and Ifit2**^-/-^** mice were harvested at days 3, 5, and 7 p.i. for flow cytometric analysis. Dot plots are representative for stains at days 3, 5 and 7 p.i. gated on total live cells and showing CD45**^int^** and CD45**^hi^** cells panel A, CD4 and CD8 cells gated on CD45**^hi^** cells depicted in Panel D and Ly6G expressing cells in the CD45**^hi^** gate panel I. Purple color in graphical representation denotes WT and green color denotes Ifit2**^-/-^** mice. Absolute numbers of CD45**^hi^**, CD45**^hi^**CD4**^+^**, CD45**^hi^**CD8**^+^** and CD45**^hi^**Ly6G**^+^** cells recovered from WT and Ifit2**^-/-^** spinal cords at indicated timepoints are compared within each group across timepoints (B, E, G, J) and between WT and Ifit2**^-/-^** mice at each timepoint (C, F, H, K) as indicated. The data were pooled from three independent experiments with N=5-10. Each dot represents a single animal. Asterix (*) represents differences that are statistically significant by Student’s unpaired t-test analysis. (*P<0.05, **P<0.01, ****P<0.0001). The error bars represent SEM

### Ifit2 deficiency impaired CD4^+^ T cell activation, and IFNγ production in the Cervical Lymph Node (CLN) upon RSA59 acute infection

Cervical lymph nodes (CLN) are the initial site for T cell activation to antigens draining from the CNS. Upon activation, T cells undergo rapid proliferation and differentiate into effectors capable of migrating to sites of infection and producing antimicrobial lymphokines[18]. IFNγ secreted by T cells plays a critical role in amplifying antigen presentation and antigen recognition via cognate T-cell-APC interaction[19]. To trace the cause of diminished CD4**^+^** T cell infiltration into the CNS on day 7 p.i, we assessed activation of CD4**^+^** T-cells in the CLN of RSA59 infected WT and Ifit2**^-/-^** mice by flow cytometric analysis of activation markers and intracellular staining for IFNγ following nonspecific PMA/IO stimulation ex vivo.

CLN of RSA59 infected Ifit2**^-/-^** mice showed reduced population of CD4**^+^** T cells (Fig. 3A-C). Activation of T cells is demarcated by specific molecular signatures including CD44 and CD62L expression[20]. Overall, naïve T cells are characterized by their CD62L**^+^**CD44**^-^**, effector/effector memory cells by their CD62L**^-^**CD44**^+^**, and central memory cells by their CD62L**^+^**CD44**^+^** phenotypes, respectively. Reduced numbers of effector/effector memory CD4**^+^** T and CD8**^+^** T cells in Ifit2**^-/-^** compared to WT mice indicate that Ifit2 deficiency causes impaired activation of T cells (Fig. 3H-N). Reduced CD4**^+^** T activation and differentiation is supported by low production of IFNγ by Ifit2**^-/-^** CD4**^+^** T cells following in vitro stimulation (Fig. 3D-G). These results suggest that Ifit2 deficiency reduces T cell activation in the CLN, which may contribute to their reduced migration to the spinal cord.

**Fig. 3:**
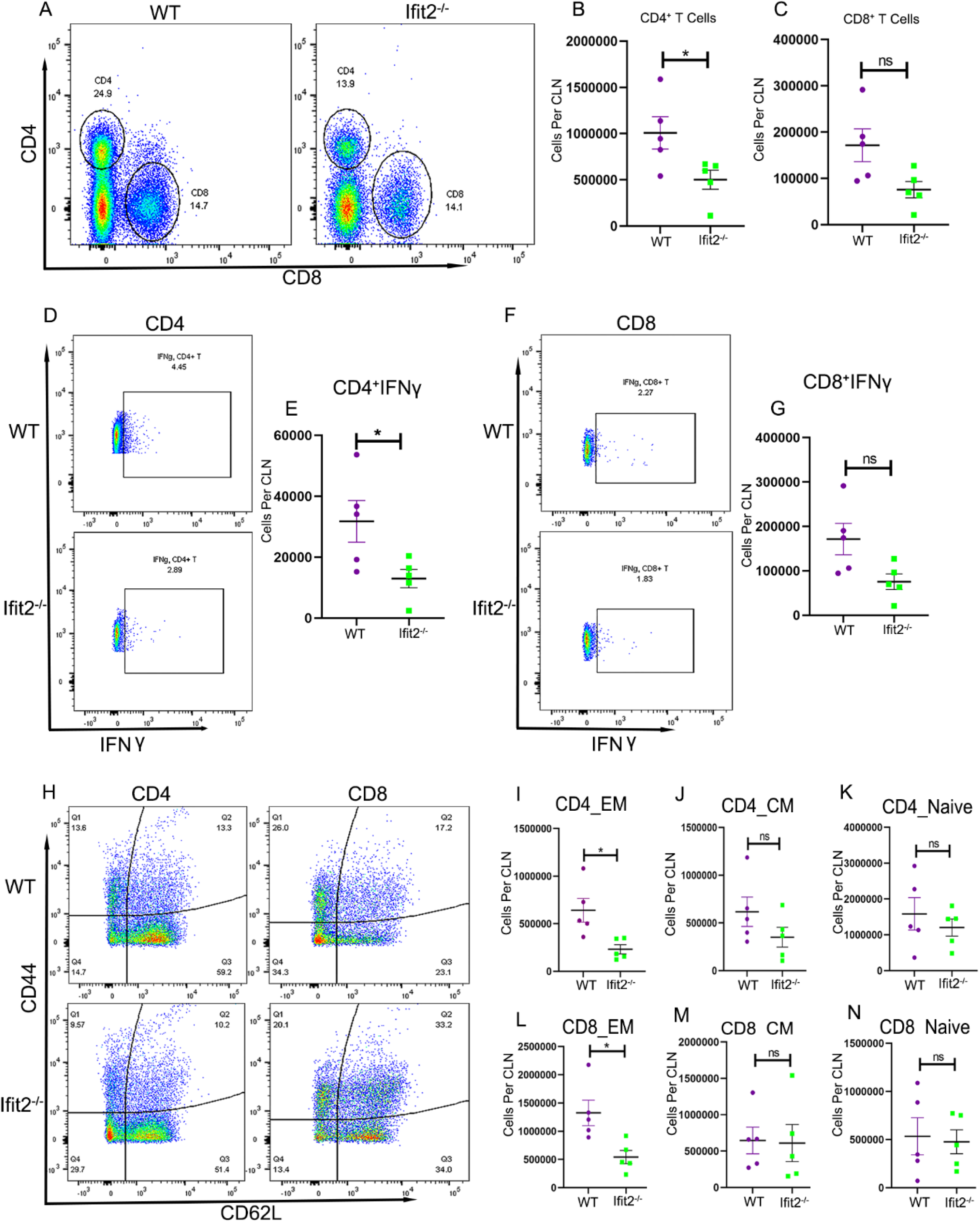
Ifit2 deficiency impairs T cell activation and IFNγ production in cervical lymph nodes (CLN) upon RSA59 acute infection. CLN derived cells from WT and Ifit2**^-/-^** RSA59 infected mice were stained for CD4, CD8, CD44, CD62L and IFNγ expression at 5 days p.i.; purple color denotes WT and green color denotes Ifit2**^-/-^** mice. IFNγ expression was assessed by intracellular staining after 6 hours in vitro stimulation with PMA/IO. (A) Representative dot plots showing gating and percentages of CD45**^hi^** CD4**^+^** and CD45**^hi^** CD8**^+^** T cells as indicated. Graphs in (B) and (C) depict absolute numbers of CD45**^hi^**CD4**^+^** and CD45**^hi^** CD8**^+^** T cells in the CLN, respectively. Dot plot in D and F show the fraction of IFNγ producing cells in CD45**^hi^** CD4**^+^** and CD45**^hi^** CD8**^+^** T cells, respectively. Panels E and G show absolute numbers of IFNγ producing CD45**^hi^**CD4**^+^** and CD45**^hi^** CD8**^+^** T cells, respectively, in CLN from individual mice. Panels H represents dot plots showing expression of CD62L and CD44 on CD4**^+^** and CD8**^+^** T cells in WT and Ifit2**^-/-^** mice. Panels I and L depict absolute numbers of CD4**^+^**CD62L**^lo^**CD44**^+^** and CD8**^+^** CD62L**^lo^**CD44**^+^** T cells in WT and Ifit2**^-/-^** mice, respectively; Panels J and M show absolute numbers of CD4**^+^**CD62L**^hi^**CD44**^+^** and CD8**^+^**CD62L**^hi^**CD44**^+^** T cells, respectively and panels K and N depict absolute number of CD4**^+^**CD62L**^hi^**CD44**^lo^** and CD8**^+^**CD62L**^hi^**CD44**^lo^** T cells, respectively. Asterix (*) represents differences that are statistically significant by Student’s unpaired t-test analysis. (*P<0.05,). The error bars represent SEM

### Ifit2 deficiency maintains tight ZO-1 staining surrounding blood vessels and maintains BBB integrity in brains of RSA59 infected mice at day 5 p.i

The blood-brain barrier (BBB) maintains control of CNS homeostasis that protects the neural tissue from toxins and pathogens. Dysregulation of these barrier functions can enhance infiltration of peripheral leukocytes into the CNS thereby promoting control of pathogens, but at the same time resulting in progression of several neurological diseases[17, 21, 22]. To assess if reduced leukocyte infiltration is associated with preserved BBB integrity, we evaluated permeability into the brain parenchyma using injection of Texas red dextran as a fluorescent dye[23]. Examination of fluorescence in the brains of infected mice following intravenous or intraperitoneal dye injection at day 5 p.i. revealed a significantly lower absolute fluorescence in the Ifit2**^-/-^** brain lysate (Fig. 4A-C). To confirm that Ifit2 deficiency is accompanied by a relatively intact BBB compared to WT mice, we also stained brain sections for ZO-1 expression, a tight junction protein abundantly found in the BBB. ZO-1 protein was abundantly expressed in Ifit2**^-/-^** mice (Fig. 4H-K) compared to WT mice (Fig. 4D-G), implying that Ifit2 contributes to loss of BBB integrity, thereby promoting CNS infiltration of peripheral immune cells. The combinatorial effects of reduced T cell priming in the CLN and tightened BBB function may thus contribute to insufficient T cell migration to mediate viral control during acute infection.

**Fig. 4:**
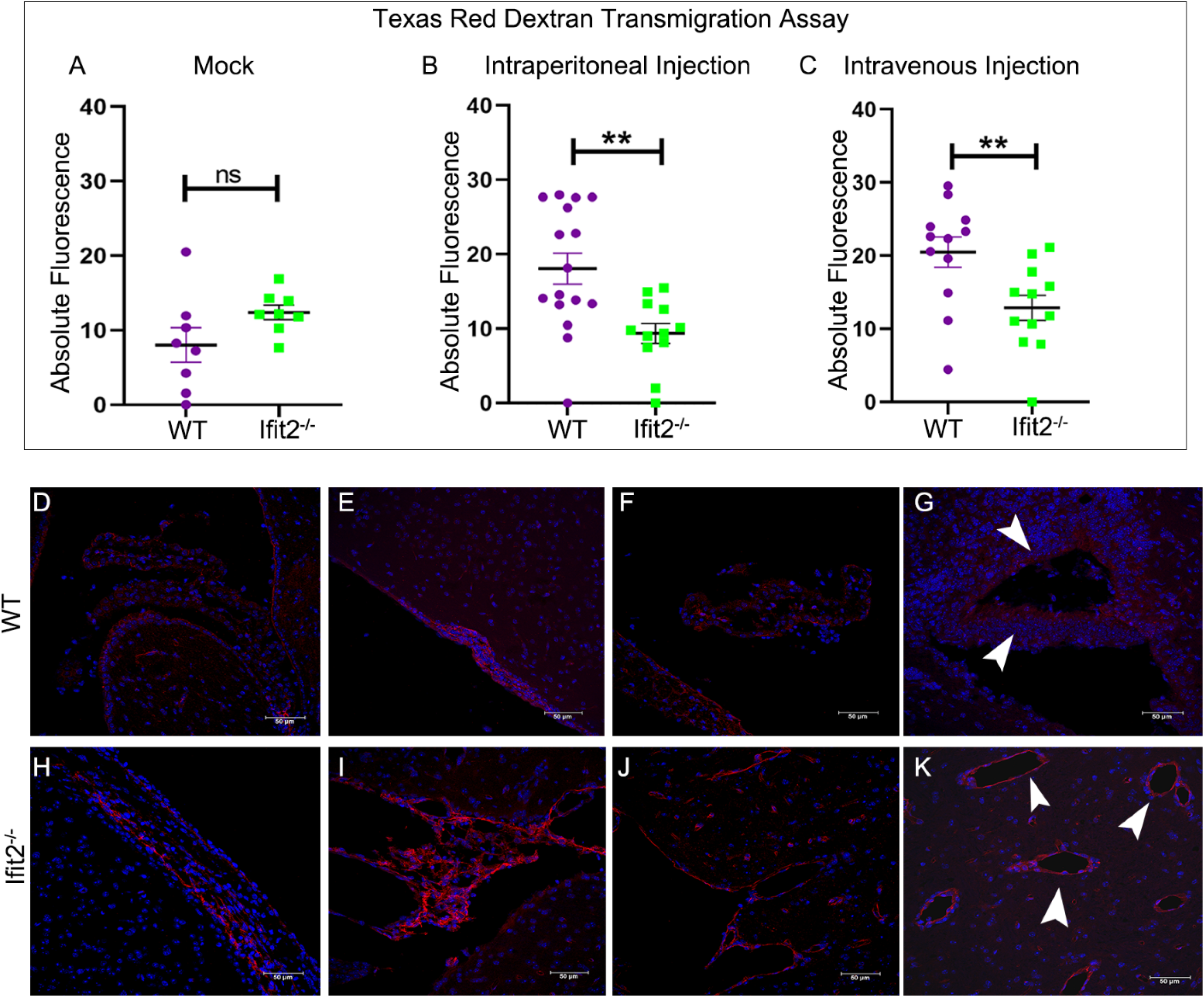
Ifit2 deficiency is associated with sustained BBB within the brains of RSA59 infected at day 5 p.i. BBB permeability was measured in infected mice following injection of Texas Red Dextran dye via the IP or IV route or mock injection of PBS at day 5 p.i. Absolute fluorescence of Texas red dextran in brain lysates was measured 15 mins post dye injection in mock infected mice (A) and RSA59 infected mice (B, C). 5-10 μm thick paraffin sections were prepared from brains of 4-5-week-old WT and Ifit2**^-/-^** RSA59 infected mice at day 5 p.i. Sections were stained for ZO-1 (red) and nuclei using DAPI (blue). Representative fluorescent images are shown from WT (Fig. 4 D-G) and Ifit2**^-/-^** mice (Fig. 4H-K). Arrowheads indicate ZO-1 staining around the blood vessels. Asterix (*) represents differences that are statistically significant by Student’s unpaired t-test analysis. (*P<0.05, **P<0.01). The error bars represent SEM.

### Ifit2^-/-^ mice exhibit severe demyelination pathology even at 500 PFU of RSA59 infection

Demyelination is the primary characteristic of the human neurological disease Multiple Sclerosis and mouse hepatitis virus (MHV) induced demyelination has provided a model to dissect inflammatory and molecular mechanisms of viral-induced demyelination. Given the high mortality of Ifit2**^-/-^** mice infected with 2000 PFUs of RSA59, we reduced the virus inoculum in Ifit2**^-/-^** mice to 500 PFUs for analysis of demyelinating pathology associated with the persistent phase of infection. The inoculum was maintained at 20,000 PFUs in WT mice as they only develop mild disease symptoms. All Ifit2**^-/-^** mice survived until day 30 p.i. with a clinical score ranging between 2-3, signified by partial to complete hind limb paralysis and severe weight loss (Fig. 5A and 5B). Spinal cords were examined by histopathological analysis at day 30 p.i. at 500 PFUs. Analysis of viral antigen by anti-N immunohistochemistry revealed readily detectable areas of viral persistence even until day 30 p.i. in all Ifit2 **^-/-^** mice, whereas WT mice showed sparse if any N staining (Fig. 5C and 5D). Viral persistence correlated with the presence of inflammatory lesions in the spinal cord indicated by H&E staining (Fig. 6A, 6B).

**Fig 5:**
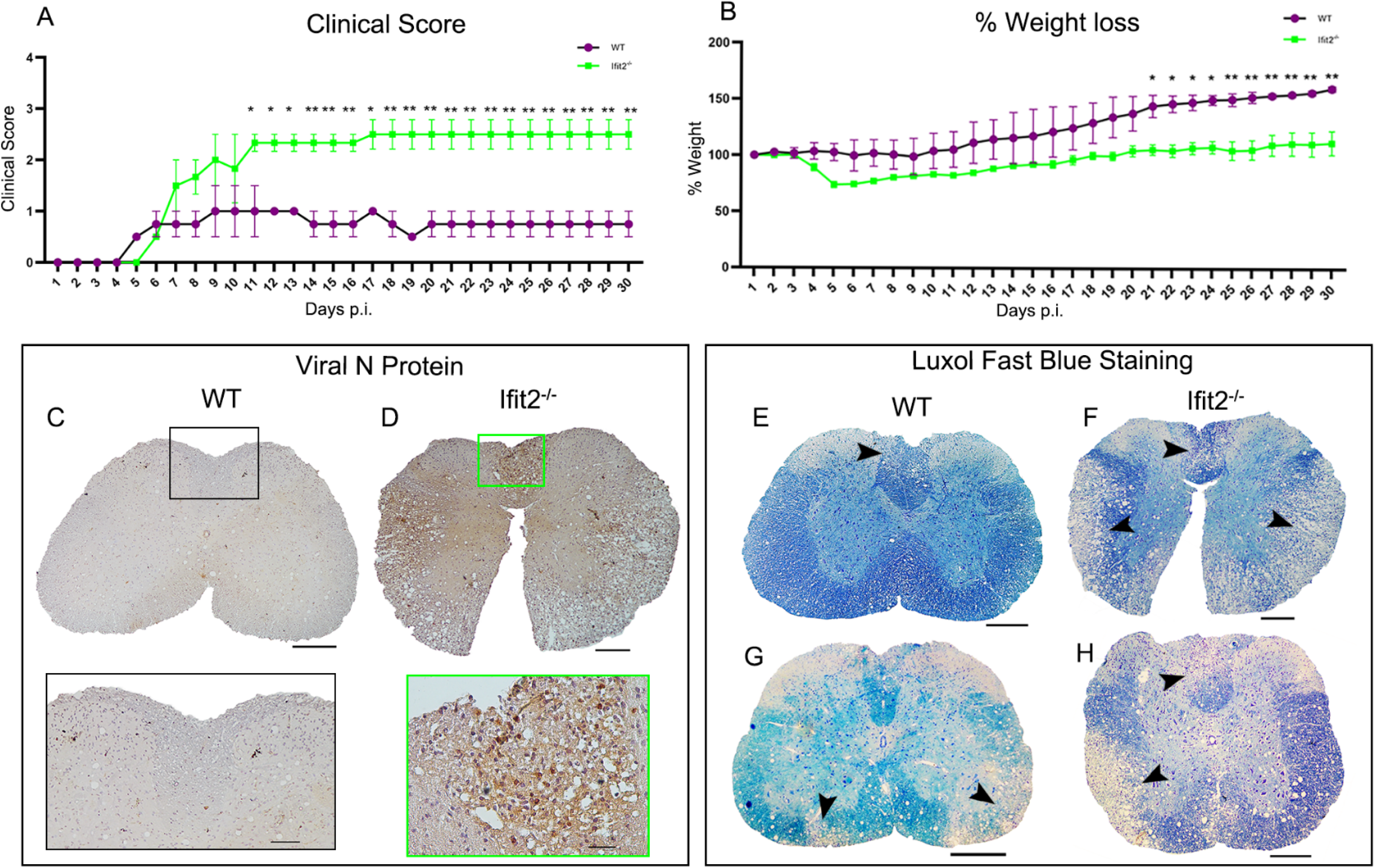
Ifit2^-/-^ mice infected with low dose RSA59 showed severe weight loss, heightened clinical score, sustained viral persistence, and severe demyelination during persistent infection. 4-5-week-old WT and Ifit2^-/-^ mice were infected intracranially with 20,000 or 500 PFU of RSA59 respectively and monitored for development of clinical disease (A) and weight loss (B). Clinical scores were assigned by an arbitrary scale of 0-4 where increasing score correlates with increased clinical impairment as described in Materials and Methods. Panels C and D show representative staining of spinal cord cross sections for viral antigen in WT and Ifit2^-/-^ mice at day 30 p.i. using anti-N antisera and corresponding enlarged region are highlighted by black square box for WT and Green for Ifit2^-/-^. Panels E-H show demyelination detected by Luxol Fast blue staining. Compared to WT mice which exhibit demyelination mainly in white matter (E, G), Ifit2^-/-^ mice show severe white matter and grey matter demyelination (F, H). Arrowheads show the region of demyelination in the spinal cord. The scale bar is 200µm for spinal cord and 50µm for the enlarged region. Asterix (*) indicates statistical significance by Two-Way ANOVA analysis for clinical score and percentage weight loss. (**P<0.01, ***P<0.001, ****P<0.0001). The error bars represent SEM.

**Fig. 6:**
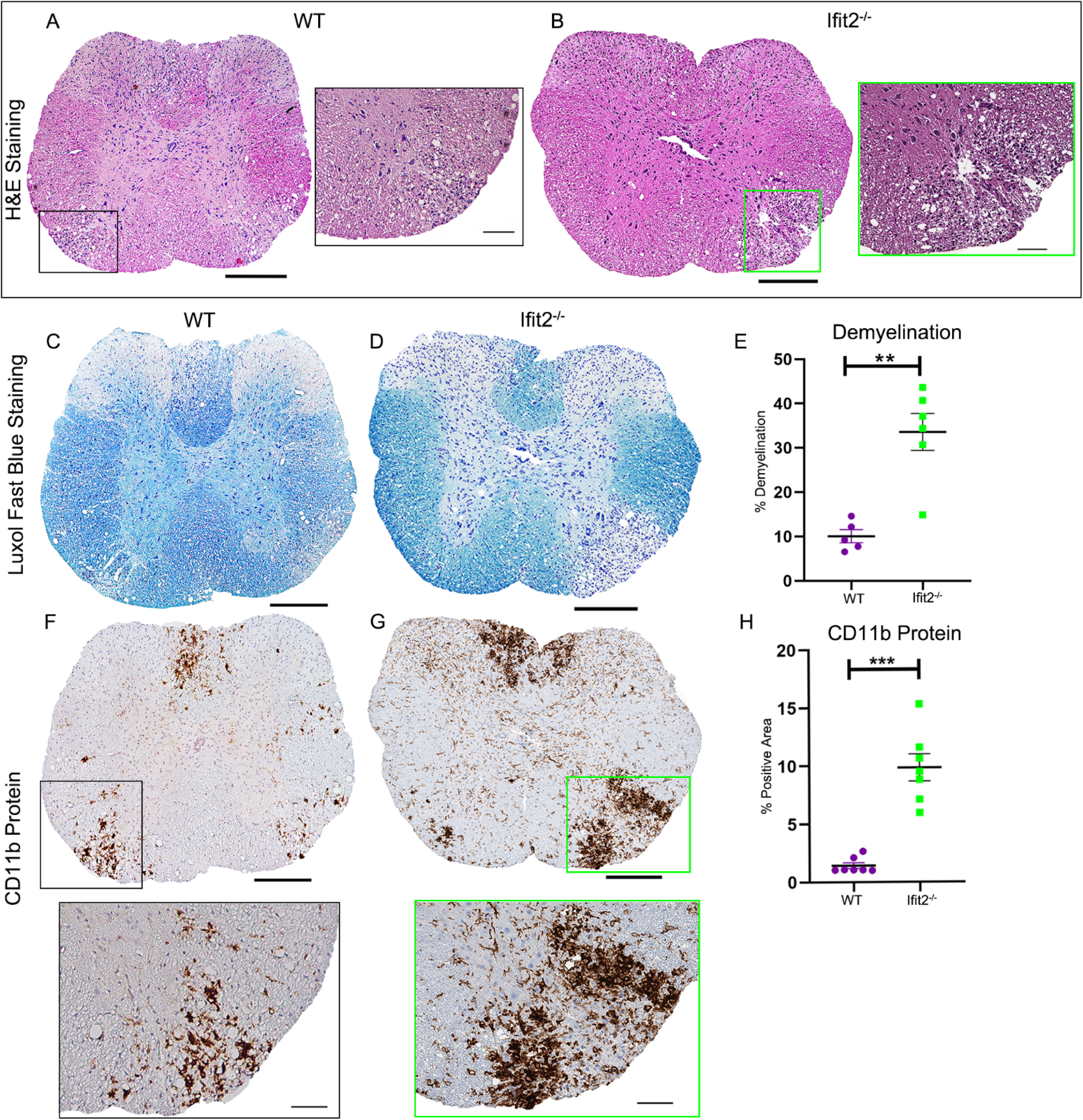
Ifit2 deficiency causes severe myelitis, chronic demyelination, and heightened accumulation of microglia/macrophages. Mice were infected as described in Fig. legend 5. Cross-sections of WT and Ifit2**^-/-^** mouse spinal cords were analyzed for the presence of inflammatory lesions by H&E (A-B), demyelination by LFB (C, D), and CD11b+ microglia/macrophages (F, G). Corresponding enlarged region is highlighted by the black square box for WT and the green box for Ifit2**^-/-^** mice. Panels E and H show quantification of white matter demyelination and microglia/macrophage activation by CD11b expression, respectively. The scale bar for spinal cord sections is 200µm and 50µm for the enlarged region. Statistical significance was calculated by unpaired Student’s t-test and Welch correction. (*P<0.05, **P<0.01, ****P<0.0001). The data represent the results from 4 or 5 independent biological experiments. The error bars represent SEM.

Luxol fast blue (LFB) staining revealed severe myelin loss in white matter of Ifit2**^-/-^** mice (Fig. 5F,5H, 6D,6E) compared to WT mice (Fig 5E, 5G, 6C, 6E). Moreover, significant grey matter pallor was evident in Ifit2**^-/-^**, but not WT spinal cords by day 30 p.i. Corresponding sections stained for CD11b showed that inflammation was primarily resolved in the grey matter of WT mice spinal cords with only few activated/phagocytic microglia/macrophages present in white matter demyelinated lesions (Fig 6F). In contrast, Ifit2**^-/-^** mice displayed a significantly large number of CD11b immunoreactive cells in both the spinal cord grey and white matter (Fig. 6G,6H). Overall, these results indicate that viral persistence is associated with the presence of morphologically defined phagocytic microglia/macrophages.

## Discussion

Type I IFNs are an integral part of the innate anti-viral immune response. IFNs exert their actions by inducing the transcription and translation of a set of genes called Interferon-induced genes (ISGs) among which ISG54/Ifit2 plays a crucial role in countering viral replication and dissemination throughout the CNS and peripheral organs[4, 8]. Ifit2 contains tetratricopeptide repeats in its structure that facilitate binding with other cellular/viral proteins as well as RNA molecules[24, 25]. Ifit2 is associated with various functions including antiviral activity, anti-tumor effects, cell migration and proapoptotic functions[26–31]. Its ability to associate with microtubules further implicates regulation of microtubule dynamics, cell proliferation, and virion assembly/transport[32]. These pleiotropic effects of Ifit2 have made it difficult to elucidate mechanisms underlying its antiviral and protective role *in vivo*.

RSA59 infection of Ifit2**^-/-^** mice is associated with enhanced viral load, impaired microglial activation, and restricted CD4**^+^** T cell migration into the brain, despite largely unaltered cytokine and chemokine production. However, the underlying mechanisms remain to be answered[11]. Two main checkpoints regulate lymphocyte migration into the CNS, the priming and activation of T cells in secondary lymphoid organs and the BBB[33, 34]. Our study revealed that the CLN of infected Ifit2**^-/-^** mice have significantly reduced numbers of CD4**^+^** T cells compared to WT mice. Also, the number of IFNγ producing, stimulated CD4**^+^** T cells is significantly reduced, supporting impaired virus-specific adaptive immune responses. CD4**^+^** T cells enhance CD8 T cells as well as humoral immunity, and protect from the development of severe MHV induced demyelination in mice[35, 36]. Impaired IFNγ production in the CNS due to reduced CD4**^+^**T cell activation may thus contribute to ineffective viral control. Reduced activation of CD4**^+^** T cells in CLN of infected Ifit2**^-/-^** mice is supported by a significant reduction in effector/effector memory CD62L**^lo^**CD44**^+^** T cells upon RSA59 infection of Ifit2**^-/-^** mice, which implicates Ifit2 in directly or indirectly affecting the maturation and activation of effector T cells in CLN. Not only does CD44 initiate T cell migration, but it is also involved in the interaction between T cells and APCs, which further promotes T cell activation[37]. A lack of CD44 expression on T cells leads to inefficient T cell migration into the CNS, and hence Ifit2**^-/-^** mice are unable to combat RSA59 virus infection, even after lowering the virus dose to 1/10th of the half of LD50 dose of RSA59. Impaired CD4**^+^**T cell activity may further be attributed to a dysfunction in CD40L-CD40 interactions. Activated CD4**^+^**T cells highly express CD40L, the ligand for CD40 expressed by microglia/monocyte /macrophages. CD40-CD40L interactions regulate cellular and humoral immune responses during viral as well as autoimmune diseases, including neurodegenerative diseases such as MS and Alzheimer’s disease[38]. Recent studies using CD40L**^-/-^** mice revealed that impaired CD4**^+^**T cell and microglia/macrophage communication causes viral persistence in the central nervous system, which in turn drives sustained activation of phagocytic microglia/macrophages and severe demyelination[20]. In addition to impaired T cell priming the integrity of the BBB, the physiological barrier between the peripheral circulation and the CNS, may contribute to a paucity of lymphocyte accumulation in the absence of Ifit2. The BBB is composed of capillary endothelial cells, astrocyte foot processes, and pericytes. The endothelial cells in the brain capillaries form tight cell-to-cell junctions composed of tight junction proteins such as claudins which interact with ZO-1,2,3. The permeability of the BBB is compromised in injury whether from trauma, multiple sclerosis, HIV infection, brain tumor, or other non-infectious inflammatory processes[21, 22]. Our data surprisingly revealed that the BBB integrity remained largely intact in RSA59 infected Ifit2**^-/-^** compared to WT mice using a Texas Red Dextran dye transfer assay and anti ZO-1 immunofluorescence. These data suggested that Ifit2 induction directly in endothelial cells or indirectly in microglia, known to interact with endothelial cells, contributes to dysregulation of tight junction proteins. Both impaired T cell priming and an intact BBB could thus contribute to the exacerbated disease progress in Ifit2 deficient mice.

An interesting aspect of Ifit2 deficiency is indeed the impaired activation of microglia, accompanied by restricted expression of CX3CR1 expression on their surface, despite a substantially high viral load[11]. Microglia are key players in maintaining CNS homeostasis[39]. Our current study demonstrated that Ifit2 deficiency impaired the activation of microglia/macrophages in the spinal cord during the acute phase of RSA59 induced neuroinflammation, similar to findings in the brain[11]. However, these microglia/macrophages remained highly phagocytic in spinal cords of Ifit2**^-/-^** mice at the chronic phase of the disease presumably due to prolonged and elevated virus persistence. Phagocytic microglia/macrophages coincident with aggravated demyelination not only in white matter, but also in the grey matter. Severe clinical disease was manifested by hind-limb paralysis. This suggested enhanced axonal damage and absence of remyelination. In this context it is important to note that microglia mediated clearance of damaged myelin is essential to promote remyelination[40]. The observation that myelin debris is clearly removed by microglia and/or infiltrated macrophages, distinct from MHV infected mice depleted in microglia, implies Ifit2 deficiency also impacts upon re-myelination. Thus, while prolonged and elevated viral persistence presumably results in elevated expression of proinflammatory factors which damage myelin, clearance of myelin debris is not impaired. The apparent lack of microglia activation and motility, reflected in morphological changes in the absence of IFN induced Ifit2 impairs viral control. However, it does not affect microglia /macrophage mediated uptake and removal of myelin debris. The inability to achieve remyelination may thus reside in impaired oligodendrocyte precursor (OPC) recruitment or differentiation due to intrinsic Ifit2 deficiency or other inhibitory factors in the lesion milieu. Conditional deletion of Ifit2 in select cell types will be needed to dissect the contribution of Ifit2 in this demyelinating disease model. Overall, our study identifies a role for Ifit2 in promoting inflammation during the acute phase of viral infection by promoting activation of T cells in the CLN, triggering permeability of the BBB, and promoting leukocyte infiltration into the CNS. The lack of functional Ifit2 results in an apparent paralysis of microglial cells to initiate and promote neuroinflammation thereby impairing viral control and resulting in viral persistence and aggravated demyelinating disease.

## Materials and methods

### Virus Infection in mice

C57BL/6 mice and homozygous Ifit2**^-/-^** mice on the C57BL/6 background bred at the breeding colony of LRI Biological Resources Unit, Lerner Research Institute, Cleveland Clinic, USA as previously described[11]. All animal experiments were carried out in strict accordance with all provisions of the Animal Welfare Act, the Guide for the Care and Use of Laboratory Animals, and the PHS Policy on Humane Care and Use of Laboratory Animals. All animal experiments were performed in compliance with protocols approved by the Cleveland Clinic Institutional Animal Care and Use Committee (PHS assurance number A3047-01). All mice were housed under pathogen-free conditions at an accredited facility at the Cleveland Clinic Lerner Research Institute and used at 4 to 5 weeks of age. The hepatotropic and neurotropic recombinant, EGFP expressing strain of MHV-A59 known as RSA59 was used in the study[41]. The virus was propagated in 17Cl1 cells and plaque assayed on DBT astrocytoma cell monolayers[16]. Mice were infected intracranially in the right hemisphere with 500 PFU and 2000 PFU for Ifit2**^-/-^** mice and 2,000 and 20,000 PFU for WT mice of RSA59 diluted in endotoxin-free, filter-sterilized PBS-BSA (Dulbecco’s phosphate-buffered saline + 0.75% BSA) in a final volume of 20 µl. Age-matched mice were mock-infected with PBS-BSA or not infected and kept as non-infected control. Clinical disease severity was graded daily using the following scale as discussed[11]. 0, no disease symptoms; 1, ruffled fur; 1.5, hunched back with mild ataxia: 2, Ataxia, balance problem and hind limb weakness: 2.5 one leg completely paralyzed, motility issue but still able to move around with difficulties; 3, severe hunching/wasting/both hind limb paralysis and mobility is severely compromised; 3.5 Severe distress, complete paralysis and moribund/ dead, 4. Our study also investigated for any phenotypic or pathological symptoms between age-matched control (non-infected) and mock-infected WT and Ifit2**^-/-^** male mice, but no such significant gross phenotypic clinical symptoms or histological changes were observed.

## Histopathological and immunohistochemical analysis

### Hematoxylin/eosin staining

RSA59 infected mice from both C57BL/6 WT and Ifit2**^-/-^** groups were sacrificed at day 5 post-infection. and day 30 p.i., and were perfused transcardially with PBS followed by PBS containing 4% paraformaldehyde (PFA). All Spinal cords were collected, post-fixed in 4% PFA overnight, and embedded in paraffin. Tissues were sectioned at 5μm and stained with hematoxylin/eosin (H&E) for evaluation of inflammation[17]. Experiments were repeated four times with 4–5 mice in each group.

### Luxol Fast Blue (LFB) staining

Mice were sacrificed at day 30 post-infection. Following transcardial perfusion with PBS and 4% paraformaldehyde, spinal cords were harvested and embedded in paraffin. 5 µm thick sections of the embedded tissues were prepared and stained with LFB stain to evaluate demyelination in the spinal cord tissues, as described previously with minor modifications[14, 20].

### Immunohistochemical staining and quantification

Serial sections from the spinal cord were stained by the avidin–biotin–immunoperoxidase technique (Vector Laboratories) using 3, 3′-diaminobenzidine as substrate, and anti-Iba1 (Wako, 1:250), CD11b (AB clonal, 1:250), anti-GFAP (Sigma, 1:500), or Anti-N (kind gift from Julian Leibowitz, Texas A&M University) (1:50) as primary antibodies. Control slides from mock-infected or uninfected mice were incubated in parallel. 7-8 sections from each infected group were randomly selected from three different sets of experiments and the expression of viral antigen, GFAP and Iba1 was quantified. Briefly, whole slides were scanned in a Leica Aperio AT2 slide scanner (Leica Microsystems, GmbH, Wetzlar, Germany) at 20x magnification and analyzed in Aperio Imagescope version 10.0.36.1805 software (Aperio) and quantified. For quantification, brightfield images of spinal cord sections were analyzed using open-source software QuPath[42]. Whole tissue sections were selected as regions of interest for analysis. Areas with tissue folding, damage or out of focus tissue were then excluded by manual annotation. Stain colors were separated into respective components by RGB color vector dependent color deconvolution. Using thresholding on the deconvolved stained images, positive pixel area was then measured as percentage.

### Blood brain barrier study

RSA59 infected mice from both C57BL/6 WT and Ifit2**^-/-^** groups were sacrificed at day 5 post-infection, and were perfused transcardially with PBS followed by PBS containing 4% paraformaldehyde (PFA). Whole brains were collected, post-fixed in 4% PFA overnight and embedded in paraffin. Tissues were sectioned at 5μm and stained for ZO-1 (Invitrogen, 1:100) protein as described[43]. For dye transmigration assay, mice were injected with 100 µl per mouse of 10mM of Texas red dextran intraperitonially and intravenously. 15 min after injection, mice were anesthetized with an I.P. injection of Ketamine and Xylazine (100 mg and 5-10 mg in 0.9 % saline per kg body weight respectively, 150 µL of the cocktail per 25 g mouse weight), followed by transcardial perfusion with PBS. The brain tissue was then harvested and homogenized. After centrifugation of the samples at 10,000 g for 15 min at 4 ◦C, the supernatant was measured to obtain raw fluorescence units (RFU) in a fluorescence plate reader at excitation/emission wavelength of 595/ 625 nm[23]. Fluorescence was plotted between mock infected WT and Ifit2**^-/-^** mice and RSA59 infected WT and Ifit2**^-/-^** mice in graphs after subtraction from autofluorescence values.

### Flow Cytometry Analysis

Mice were perfused with PBS and spinal cord were homogenized in 4 ml of Dulbecco’s PBS (pH 7.4) using Tenbroeck tissue homogenizers. Following centrifugation at 450 g for 10 min, cell pellets were resuspended in RPMI containing 25 mM HEPES (pH 7.2), adjusted to 30% Percoll (Sigma) and underlaid with 1 ml of 70% Percoll. Following centrifugation at 800 g for 30 minutes at 4°C, cells were recovered from the 30%-70% interface, washed with RPMI, and suspended in FACS buffer (0.5% bovine serum albumin in Dulbecco’s PBS). Also, deep cervical lymph nodes were harvested, homogenized in 4 ml RPMI containing 25 mM HEPES (pH 7.2), passed through 70 μm filters followed by 30 μm filters to obtain single-cell suspensions. Following centrifugation at 45 g for 10 min, cell pellets were resuspended in FACS buffer. For intracellular staining, CNS-derived cells were stimulated for 6 h with phorbol 12-myristate 13-acetate (PMA) (10 ng/ml) (Acros Organics, Geel, Belgium) and ionomycin (1 μM) (Calbiochem, Spring Valley, CA, USA), with Monensin (2 μM) (Calbiochem) added for the last 2 h. Following stimulation, surface molecules were detected as described below. Cells were permeabilized using Cytofix/Cytoperm solution (BD Biosciences) and incubated for 30 min on ice with fluorescent monoclonal antibody (mAb) specific for IFN-γ (XMG1.2; BD Biosciences). Cells were then washed using Perm/Wash buffer according to the manufacturer’s instructions[44]. Cells were counted using an automated cell counter (Invitrogen) to obtain the numbers of total leukocytes. 1 million cells were stained for flow cytometry[11, 20]. Fc receptors were blocked with 1% polyclonal mouse serum and 1% rat anti-mouse CD16/ CD32 (clone 2.4G2; BD Biosciences, San Jose, CA) monoclonal antibody (MAb) for 20 minutes. Specific cell types were identified by staining with fluorescein isothiocyanate (FITC)-, phycoerythrin (PE)-, peridinin chlorophyll protein (PerCP)-, or allophycocyanin (APC)- conjugated MAb for 30 minutes on ice in FACS buffer. Expression of surface markers was characterized with MAb (all from BD Biosciences except where otherwise indicated) specific for CD45 (clone Ly-5), CD4 (clone GK1.5), CD8 (clone 53-6.7), CD11b (clone M1/70), Ly- 6G (clone 1A8), and NK1.1 (clone PK136), CD44 (clone IM7), CD62L (MEL-14), IFNγ (clone XMG1.2). Samples were analyzed using a BD LSRFortesa flow cytometer (BD Biosciences) and FlowJo 10 software (Treestar, Inc., Ashland, OR). First, doublet exclusion using FSC-A and FSC-W was performed, and then cells were gated based on forward scatter (FSC), and side scatter (SSC) to focus on live cells. Cells were gated from a primary gating on CD45. Single colors and FMOs were used in all the experiments.

### Statistical Analysis

All immunohistochemical analysis and flow cytometry statistical analysis was performed by Student’s unpaired t-test. Data were analyzed using Prism software (GraphPad Prism 8). Two-Way ANOVA analysis for clinical score and weight loss. (**P<0.01, ***P<0.001, ****P<0.0001)

## Acknowledgement

We thank the Lerner Research Institute, Cleveland Clinic where all the experiments were done. We thank the LRI Biological Resources Unit for assistance with animal care and handling. The authors also thank the members of the LRI Imaging Core: Diane Mahovlic and Andrelie Branicky for histology services, Kelly Simmerman for Immunohistochemistry, Dr. Gauravi Deshpande for help with imaging and analyses, and Dr. Judy Drazba for project consultation. We thank CSIR for providing fellowship to MS, IISER-Kolkata for providing fellowship to DC. We also want to thank SERB-POWER (Promoting Opportunities for Women in Exploratory Research) program for providing research funds to JDS for structured effort toward enhanced diversity in research to ensure equal access and weighted opportunities for Indian women scientists engaged in research and development activities

## Authors Contribution

Conceptualization: Jayasri Das Sarma

Data curation: Madhav Sharma, Debanjana Chakravarty, Amy Burrows, Patricia Rayman, Jayasri Das Sarma

Formal analysis: Madhav Sharma, Debanjana Chakravarty, Ajay Zalavadia, Nikhil Sharma, Amy Burrows, Patricia Rayman, Jayasri Das Sarma

Funding Acquisition: Jayasri Das Sarma, Ganes C. Sen Investigation: Jayasri Das Sarma

Methodology: Madhav Sharma, Debanjana Chakravarty, Amy Burrows, Patricia Rayman, Jayasri Das Sarma

Project administration: Jayasri Das Sarma Resources: Jayasri Das Sarma, Ganes C. Sen

Software: Madhav Sharma, Debanjana Chakravarty, Ajay Zalavadia, Jayasri Das Sarma.

Supervision: Jayasri Das Sarma, Cornelia Bergmann, Ganes C. Sen. Validation: Jayasri Das Sarma, Cornelia Bergmann, Ganes C. Sen. Visualization: Jayasri Das Sarma.

Writing – original draft: Madhav Sharma, Debanjana Chakravarty, Jayasri Das Sarma. Writing – review & editing: Jayasri Das Sarma, Lawrence C. Kenyon, Cornelia Bergmann, Ganes C. Sen.

